# Maintaining the methionine residues of the chaperone Spy in a reduced state is crucial for periplasmic proteostasis

**DOI:** 10.1101/2024.12.12.628106

**Authors:** Laurent Loiseau, Nathan De Visch, Alexandra Vergnes, Jean Armengaud, Maxence S. Vincent, Benjamin Ezraty

## Abstract

The bacterial cell envelope is exposed to various stresses, including oxidative stress caused by different types of oxidants, such as reactive oxygen species (ROS) and reactive chlorine species (RCS). In *E. coli*, the reduction of chlorate into chlorite, a toxic RCS compound, induces the expression of the MsrPQ system, which repairs periplasmic proteins oxidized at methionine residues (methionine sulfoxide, Met-O). In this study, using a proteomic-based approach, we show that chlorite stress also triggers the overproduction of the periplasmic chaperone Spy. This response is mediated by the activation of the BaeSR two-component system. Furthermore, both *in vivo* and *in vitro* evidence reveal that Spy’s susceptibility to oxidation is critical for its chaperone activity. We demonstrate that the MsrPQ repair system ensures Spy’s functionality by reducing its Met-O, thereby safeguarding its role in periplasmic protein homeostasis. Overall, this work reveals Spy as a key target of chlorite-induced oxidative damage and underscores the essential role of MsrPQ in preserving periplasmic protein quality control.

## INTRODUCTION

Preserving protein homeostasis (proteostasis) and ensuring protein functionality are crucial processes for cells [1]. In bacteria, the cell envelope in general and the periplasm of Gram-negative bacteria in particular, are exposed to endogenous and exogenous oxidative stresses that damage proteins [2,3]. The accumulation of damaged proteins disrupts important biological processes and can result in cellular dysfunction and even cell death. To address this damage and maintain proteostasis, bacteria rely on a combination of oxidative species-scavenging enzymes, chaperones, proteases, and repair enzymes [4,5].

In the periplasm, where ATP is absent, folding and degradation factors operate through ATP-independent mechanisms. In *E. coli*, a network of periplasmic chaperones and proteases has been identified [5,6], including the protease/chaperone DegP, the prolyl isomerase FkpA, the chaperones SurA and Skp, which assist in outer membrane insertion, and the chaperone Spy (Spheroplast Protein Y). Spy is a 16 kDa chaperone that prevents protein aggregation and promotes client protein refolding at sub-stoichiometric concentrations [7]. Spy’s active form is a cradle-shaped dimer containing a large hydrophobic surface at its core and basic residues on its edges [8,9]. The immunity protein 7 (Im7) serves as a model client for Spy [8], enabling investigation of the mechanism of chaperone action and demonstrating that Spy uses complementary charge interactions to bind to its unfolded substrates [8,10]. The expression of the *spy* gene is regulated by the Cpx and Bae envelope stress pathways [7], resulting in its upregulation during stress conditions such as exposure to ethanol, copper, and tannins [7,11].

The periplasm is more oxidizing than the cytoplasm and is the first compartment to encounter exogenous oxidizing agents. Consequently, reduction enzymes targeting the highly susceptible sulfur-containing amino acids cysteine (Cys) and methionine (Met) - specifically the Dsb (disulfide bond) and Msr (methionine sulfoxide reductase) enzyme families - are essential components of periplasmic protein quality control [3]. MsrPQ, the periplasmic Msr system widely conserved in Gram-negative bacteria, reduces protein-bound Met-O [12]. In *E. coli*, the expression of the *msrPQ* operon is induced by reactive chlorine species (RCS), such as HOCl (bleach), *N*-chlorotaurine and chlorite (ClO_2_⁻), via the HprSR pathway [12–14]. The toxic oxidizing agent chlorite, produced by the reduction of chlorate (ClO₃⁻) by nitrate reductases, oxidizes Met in periplasmic proteins, supporting the role of MsrPQ as an anti-chlorite defense system [14].

Here, we identify the Spy chaperone as a key player during chlorite treatment. The Bae pathway appears to be activated by chlorite stress, leading to the overproduction of Spy. We found that Spy is oxidized under chlorite stress, which is reversible through the action of MsrPQ. Further analyses revealed that the oxidation of three Met residues in the cradle region disrupts Spy’s chaperone activity. Overall, we propose a model in which MsrPQ plays a crucial role in preserving chaperone function in the periplasm, highlighting its importance in periplasmic proteostasis.

## RESULTS

### Exposure to chlorite induces Spy synthesis

In a previous study [14], we found that chlorite induces the expression of the *hiuH-msrPQ* operon. We investigated whether other proteins were overproduced in *E. coli* during chlorite stress. To this end, we performed a global label-free proteomic analysis of cells, recording 798,574 MS/MS spectra, identifying 15,818 peptide sequences, and monitoring 1,496 proteins. Comparative proteomics revealed that Spy, a periplasmic chaperone, was the most overproduced protein in response to chlorite, with a fold change of 15.3 (Fig. 1A and Supplementary Table 1). Consistent with our previous findings, the top five proteins with increased abundance included HiuH and MsrP, ranked second and fourth, respectively.

**Figure 1:**
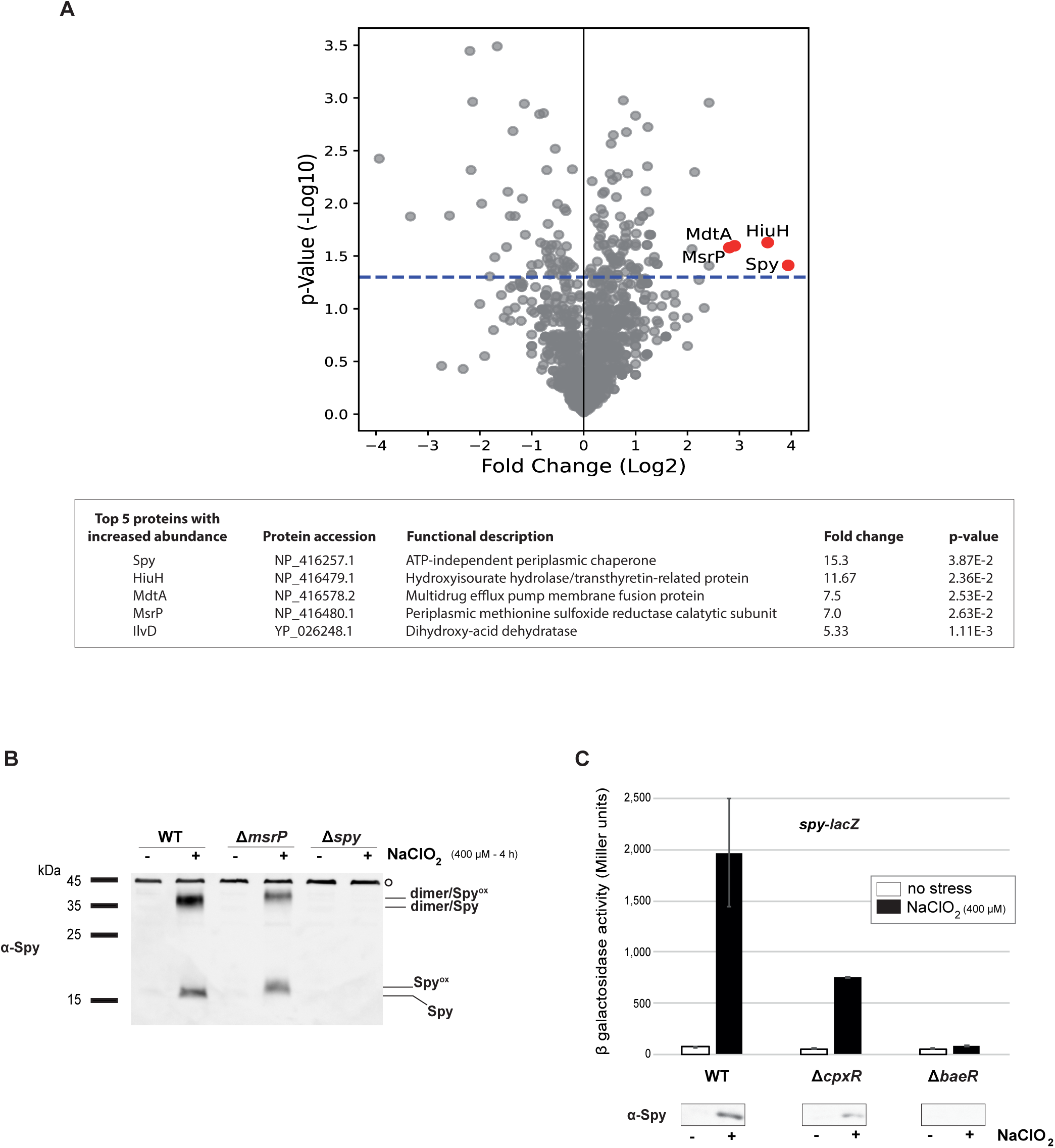
Chlorite exposure induces Spy production and oxidation. **A.** Volcano plot of mass spectrometry results showing proteins significantly upregulated by chlorite exposure. The four proteins with the greatest fold changes are highlighted in red. The horizontal dashed line indicates the p-value threshold. A table summarizing the accession numbers, functional descriptions, fold changes, and p-values for the top five proteins is provided. **B.** Immunodetection of Spy in the periplasmic fraction of WT, Δ*msrP* (CH380), and Δ*spy* (LL1392) cells. Spy levels were detected using an anti-Spy antibody. The oxidized form of Spy (Spy^ox^), identified by a mobility shift on SDS-PAGE, is indicated. Molecular weights are indicated. A non-specific band is indicated by open circle. **C.** *spy* expression in response to chlorite exposure. β-galactosidase assays were perfomed on WT (LL1322), Δ*cpxR* (LL1324), and Δ*baeR* (LL1378) strains are shown, with error bars representing standard deviations (n=3). Spy protein levels were also confirmed by western blotting of whole-cell extracts from WT, Δ*cpxR* (LL1198), and Δ*baeR* (LL1538) strains using an anti-Spy antibody.

To further investigate Spy production under chlorite stress, we analyzed the periplasmic protein profile of wild-type *E. coli* cultures exposed to chlorite or left untreated. Immunoblotting revealed a massive accumulation of Spy after chlorite exposure in the wild-type strain, but no signal was detected without chlorite treatment or in a Δ*spy* mutant strain (Fig. 1B). Interestingly, we also observed the Spy dimer, even under denaturing gel conditions (Fig. 1B). We further confirmed the induction of *spy* in response to chlorite exposure using a *spy-lacZ* transcriptional fusion that exhibits an approximately thirtyfold increase under chlorite stress conditions (Fig. 1C). Together, these results indicate that Spy is among the most overproduced proteins in response to chlorite stress.

### The BaeRS signaling pathway modulates the expression of *spy* in response to chlorite stress

In *E. coli*, *spy* expression is regulated by the CpxRA and BaeRS envelope stress response pathways [11]. To identify the pathway involved during chlorite stress, we quantified *spy-lacZ* expression in different genetic backgrounds (Fig. 1B). In *ΔcpxR* cells, *spy* expression increased under chlorite treatment but remained lower than in wild-type cells, whereas no increase was observed in *ΔbaeR* cells. (Fig. 1B). Additionally, Spy production during chlorite stress was significantly reduced in Δ*baeR* cells compared to wild-type and Δ*cpxR* cells (Fig. 1B). These observations indicate that *spy* expression following chlorite treatment is regulated by the BaeRS system rather than by CpxRA (Fig. 1B). The role of BaeRS in response to chlorite stress was also confirmed by the global proteomic analysis, which identified several proteins, such as MdtA and MdtB, encoded by Bae-regulated genes among the most abundant under chlorite stress (Fig. 1A and Supplementary Table 1) [15].

### Spy is specifically induced by chlorite

To determine whether *spy* induction is specific to chlorite or can be triggered by other ROS/RCS, *spy-lacZ* expression was analyzed following exposure to sublethal concentrations of various oxidants, including H_2_O_2_, paraquat (a superoxide generator), diamide, HOCl, and *N*-chlorotaurine. Tannic acid, a known inducer of *spy* expression [7], was used as a positive control. Among these oxidants, only chlorite significantly increased *spy* expression, while all others caused negligible changes (Fig. 2A).

**Figure 2:**
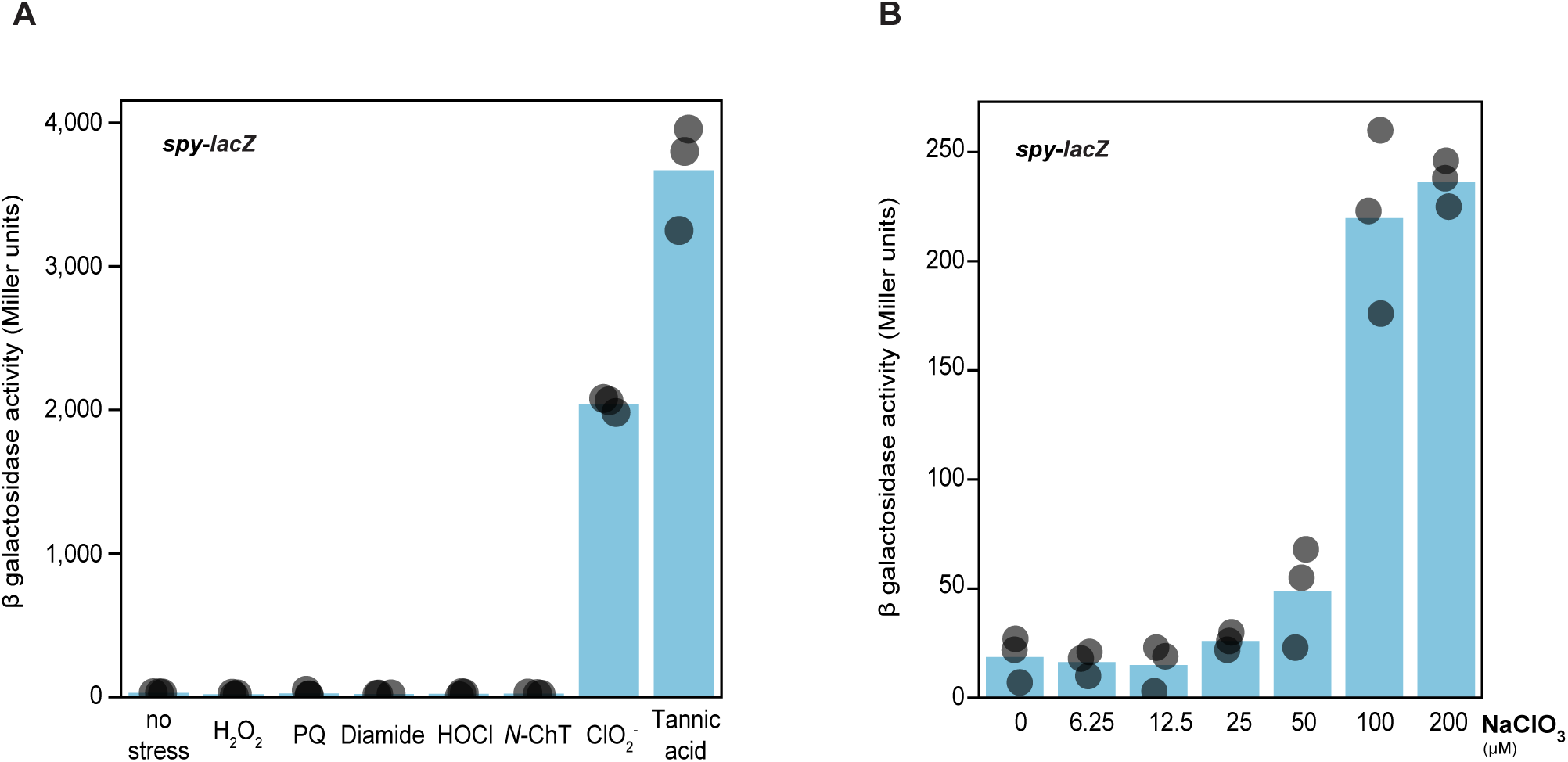
Spy induction is specific to chlorite stress. *spy* expression in response to various oxidants. **A.** β-galactosidase assays of WT (LL1322) treated aerobically with 5 mM H_2_O_2_, 0.3 mM paraquat, 0.3 mM diamide, 4 mM HOCl, 1 mM *N*-ChT, 0.4 mM NaClO_2_, or 0.25 mM tannic acid for 4 hours. **B.** β-galactosidase assays were performed on WT (LL1322) treated anaerobically overnight with NaClO₃ (6.25–200 µM). Error bars indicate standard deviations (n=3).

Given that chlorite is produced via chlorate reduction by nitrate reductases under anaerobic conditions [16–18], we investigated whether chlorate induces *spy* expression. Adding 50 to 200 µM of chlorate to LB medium led to a notable increase in β-galactosidase activity from the *spy-lacZ* fusion (Fig. 2B). These results demonstrate that *spy* is also upregulated under anaerobic conditions in the presence of chlorate.

### MsrP repairs chlorite-oxidized Spy

The deletion of *msrP* alters Spy migration during electrophoresis under denaturing conditions (Fig. 1B), causing a mobility shift characteristic of Met-O-containing polypeptides [19]. This suggests that (i) chlorite induces both Spy expression and its oxidation and (ii) that MsrP is involved in the redox control of Spy. To test this hypothesis, we monitored the redox state of Spy in the presence or absence of MsrP. In an Δ*msrP* strain, expressing *msrP* from a plasmid restored Spy’s reduced mobility, whereas Spy remained oxidized in the strain carrying an empty vector (Fig. 3). MsrP production, induced either by chlorite treatment or plasmid expression, was confirmed by immunoblotting (Fig. 3). Altogether, these findings indicate that, during chlorite stress, (i) Spy’s Met residues are oxidized, and (ii) MsrP repairs and maintains Spy in its reduced form.

**Figure 3:**
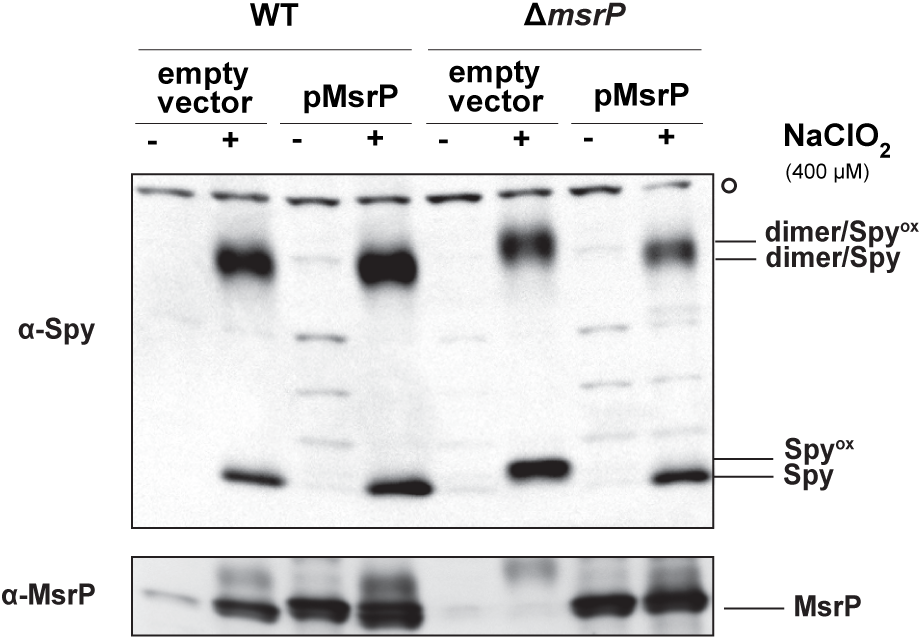
Reversible oxidation of Spy is controlled by MsrP. Spy and MsrP protein levels were analyzed by immunoblotting and detected using anti-Spy and anti-MsrP antibodies. The oxidized form of Spy (Spy^ox^), identified by a mobility shift on SDS-PAGE, is indicated. Molecular weights are indicated. A non-specific band is indicated by open circle.

### Identification of methionine residues important for Spy function

Whether oxidation of Spy’s Met is critical for its chaperone activity is unknown. Spy contains eleven Met residues, some concentrated at the N-terminus in two doublets (Met37-Met38 and Met50-Met51) while others (69, 87, 108 and 120) are located more centrally or at the C-terminus part (Fig. 4A). Quan *et al.* [7] identified these latter position as conserved among Spy homologs in enterobacteria, proteobacteria, and in some cyanobacteria. Interestingly, three of these Met residues (Met69; Met87 and Met108) are also conserved with the homologous CpxP protein from *E. coli* (Fig. 4A) [20,21], and structural analysis revealed that they are located at the core of the substrate-binding cradle (Fig. 4B). Based on this analysis, we focused on the N-terminal Met hot spot (Met37-Met38 and Met50-Met51), that we refer to as group 1, and the three core Met (Met69, Met87, and Met108) as group 2. We substituted these Met residues with either Ala (A), to test the physicochemical constraints at each position, or Gln (Q), a Met-O mimetic, to assess the consequences of oxidation at those positions [22,23].

**Figure 4:**
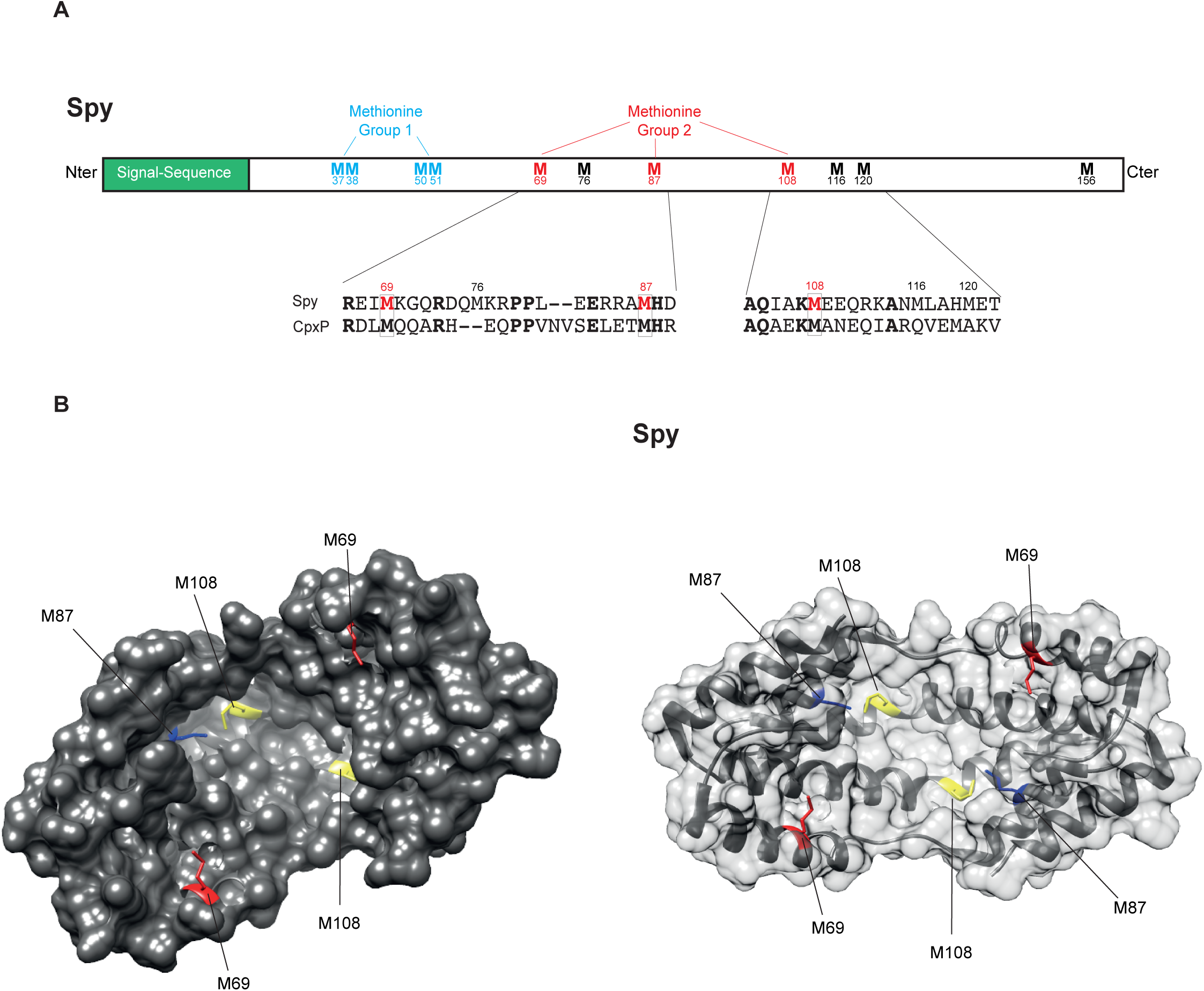
Structural features of the Spy protein. **A.** Schematic of the Spy protein indicating the N-terminal Met hotspot (Met37-Met38 and Met50-Met51, Group 1, in blue) and the core Met residues (Met69, Met87, and Met108, Group 2, in red). Conservation of these Met residues with corresponding residues in the CpxP protein is also shown. **B.** Structure of the *E. coli* Spy protein (PDB:3O39). The Spy dimer is shown in surface view, with core Met residues color-coded: Met69 in red, Met87 in blue, and Met108 in yellow.

Given that deletion of *spy* did not result in any observable phenotype under either normal growth or stress conditions, we could not test the functionality of the Met-to-Ala and Met-to-Gln variants through complementation. Instead, we performed a multicopy suppression test, utilizing the overproduction of Spy to suppress the novobiocin or clindamycin sensitivity in a Δ*skp* Δ*fkpA* mutant background (Fig. 5A) [24]. We monitored the antibiotics sensitivity of the Δ*skp* Δ*fkpA* strain, in which the *spy* gene was deleted to prevent expression of the endogenous wild-type *spy*, carrying plasmids expressing the different Spy variants. Our results show that the group 1 N-terminal Met residues are not essential for Spy’s function, as variants with up to four Met-to-Ala or Met-to-Gln substitutions suppressed antibiotic sensitivity as effectively as wild-type Spy (Fig. 5B).

**Figure 5:**
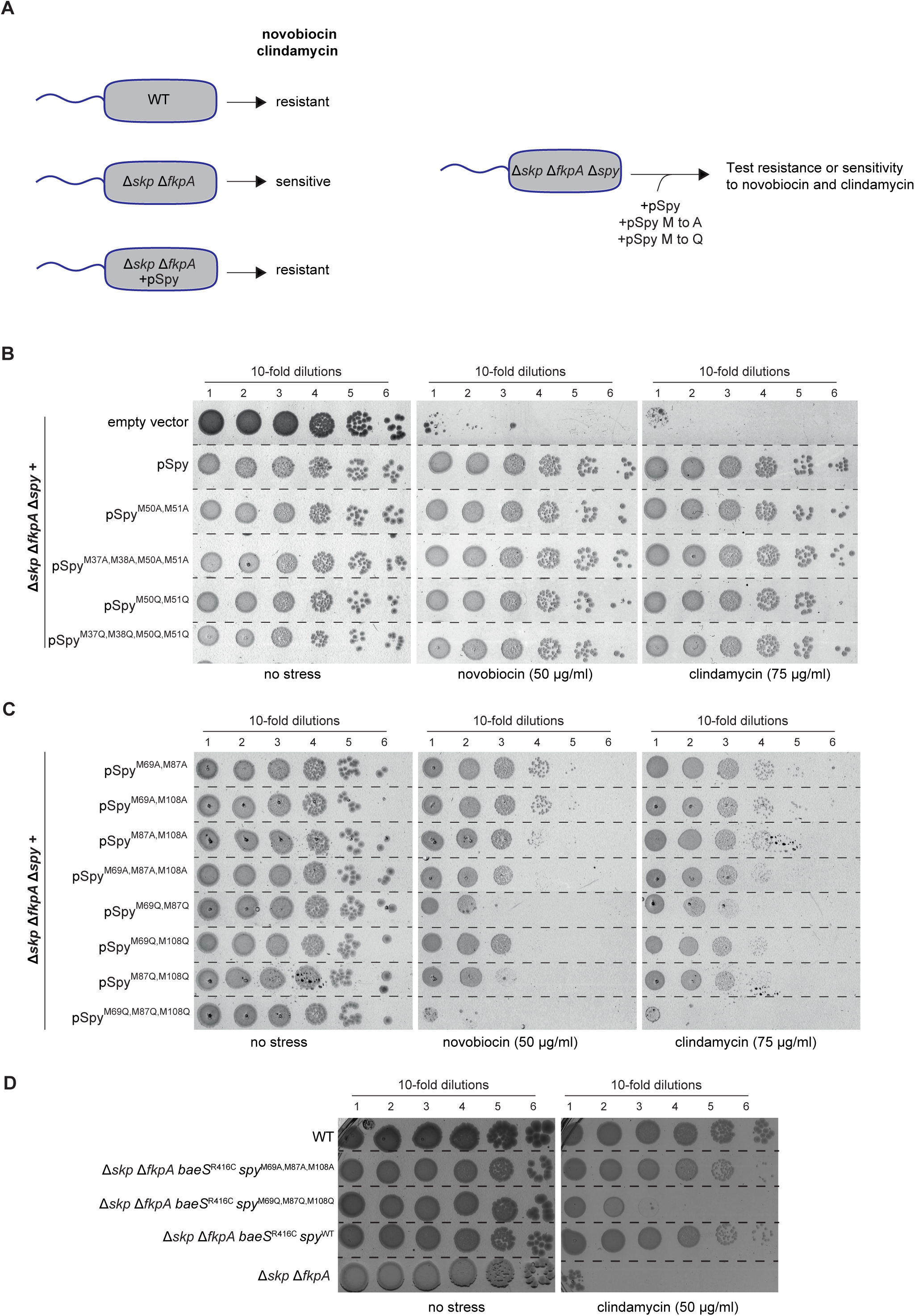
Importance of Spy’s Met residues. **A.** Unlike WT *E. coli,* the Δ*skp* Δ*fkpA* strain exhibits sensitivity to novobiocin and clindamycin. Overproduction of Spy suppresses this phenotype, providing a functional assay to evaluate Spy variants. **B.** Δ*skp* Δ*fkpA* Δ*spy* (LL1414) cells carrying an empty vector or a plasmid expressing wild-type Spy or its variants of group 1 (**B**) or group 2 (**C**) are spotted onto LB plates containing ampicillin and IPTG, with or without the addition of novobiocin (50 µg/ml) or clindamycin (75 µg/ml). **D.** WT, Δ*skp* Δ*fkpA* (LL1334), Δ*skp* Δ*fkpA baeS*^R416S^ (LL1590), Δ*skp* Δ*fkpA baeS*^R416S^ *spy*^M69Q,M87Q,M108Q^ (LL1594), Δ*skp* Δ*fkpA baeS*^R416S^ *spy*^M69A,M87A,M108A^ (LL1596) are spotted onto LB plates supplemented with or without clindamycin (50 µg/ml).

The same assay was performed for the group 2 Met (Fig. 5C). The Met-to-Ala substitution mutants showed a reduced ability to suppress antibiotic sensitivity, with a stronger effect in the triple mutant (M69A/M87A/M108A) (Fig. 5C). Thus, positions 69, 87, and 108 are functionally important, with a low tolerance for substitutions at these positions.

The Met-to-Gln mutants, mimicking Met oxidation, were unable to suppress antibiotic sensitivity (Fig. 5C). The double M69Q/M87Q and the triple M69Q/M87Q/M108Q substitutions were particularly affected (Fig. 5C).

To rule out potential effects associated with the use of plasmids, we introduced the triple Met-to-Ala and Met-to-Gln substitution alleles into the chromosome of a strain lacking *skp* and *fkpA* and carrying the R416C mutation in the *baeS* gene [7], which renders the Bae pathway constitutively active. The resulting Met-to-Gln strain exhibited sensitivity to clindamycin, whereas the Met-to-Ala strain appeared resistant, similar to the strain containing the wild-type *spy* allele (Fig. 5D).

These genetic screens identified the N-terminal Met37, Met38, Met50, and Met51 as non-essential for Spy’s function. In contrast, Met69, Met87, and Met108 are crucial and their oxidation causes loss of activity.

### Methionine oxidation in the cradle impairs Spy’s chaperone activity

To determine whether Met oxidation in the cradle region impairs Spy’s chaperone activity, we conducted *in vivo* and *in vitro* assays.

The chaperone activity of Spy can be evaluated *in vivo* by monitoring its ability to stabilize the Im7 protein, a known Spy substrate [7]. This is achieved using a stability biosensor, which incorporates the unstable protein Im7-L53A/I54A between the domains of β-lactamase. If Spy stabilizes Im7, the β-lactamase is reconstituted, rendering the strain resistant to penicillin (Fig. 6A) [7,25]. As expected, strains expressing the “Super Spy^Q100L^ Variant” exhibited enhanced resistance to penicillin V compared to cells expressing wild-type Spy (Fig. 6B) [25]. In contrast, strains lacking Spy showed sensitivity to the antibiotic (Fig. 6B). Remarkably, while the expression of the Met-to-Ala substitution mutant (Spy^M69A/M87A/M108A^) displayed an intermediate antibiotic sensitivity phenotype, we found that expressing the Met-to-Gln substitution mutant (Spy^M69Q/M87Q/M108Q^) phenocopied the absence of Spy (Fig. 6B).

**Figure 6:**
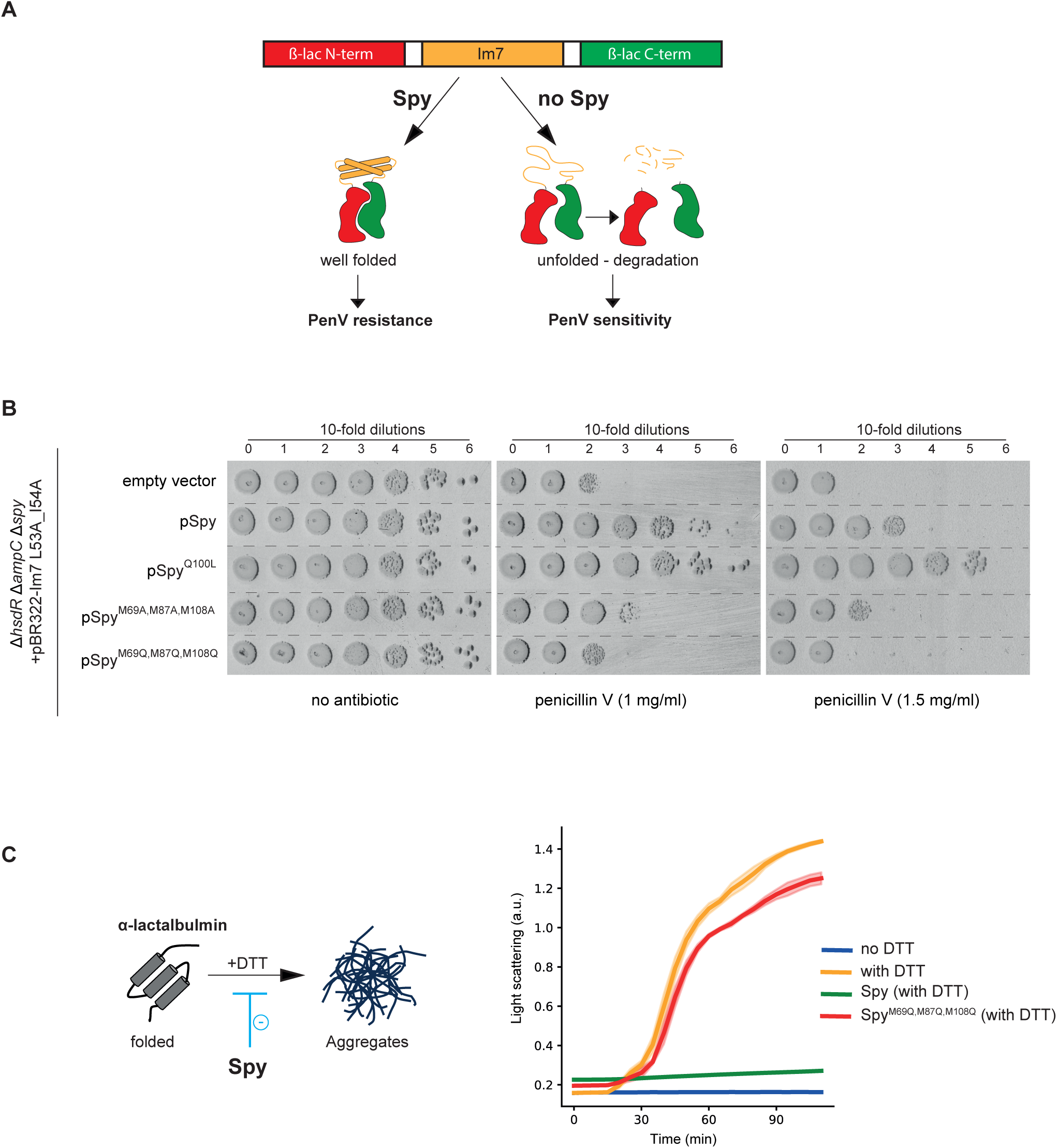
Impact of Met oxidation on Spy’s chaperone activity. **A.** The unstable Im7-L53A/I54A protein, a known substrate of Spy, has been inserted between the domains of β-lactamase. In the presence of Spy, Im7 is properly folded, and β-lactamase is reconstituted, making the strain resistant to penicillin V. In the absence of Spy, Im7 is unfolded and degraded, rendering the strain sensitive to penicillin V. **B.** Δ*hsdR* Δ*ampC* Δ*spy* (HW38) cells carrying plasmids expressing the unstable Im7-L53A/I54A protein and Spy^WT^, Spy^Q100L^, Spy^M69A,M87A,M108A^ or Spy^M69Q,M87Q,M108Q^ were grown in LB supplemented with kanamycin, tetracycline, and IPTG at 37°C until an OD_600_ of 0.6. Ten-fold serial dilutions were spotted onto LB plates containing IPTG, with or without penicillin V (1 and 1.5 mg/ml). **C.** α-lactalbumin aggregation monitored by light scattering over time without DTT (blue), with DTT (orange), with DTT and Spy^WT^ (green) and DTT and Spy^M69Q,M87Q,M108Q^ (red). The mean values are shown, with standard deviation represented by a shaded area.

To assess the chaperone activity of Spy *in vitro*, we purified Spy and its variant version and determined their ability to prevent the aggregation of dithiothreitol-reduced α-lactalbumin (α-LA) (Fig. 6C) [10,26]. We found that Spy^WT^ prevents α-LA aggregation, whereas the Spy^M69Q/M87Q/M108Q^ variant displayed no anti-aggregation capability (Fig. 6C), indicating a loss of chaperone activity. Overall, these results demonstrate that the Spy^M69Q/M87Q/M108Q^ variant is nonfunctional and that oxidation of Met 69, 87, and 108 results in a loss of Spy’s chaperone activity.

## DISCUSSION

In this study, we uncovered that the periplasmic chaperone Spy plays a pivotal role in responding to chlorite stress, a response triggered by the activation of the BaeSR stress pathway. We provided evidence that Spy is vulnerable to oxidation, particularly at three conserved Met residues—Met69, Met87, and Met108—which are critical for its chaperone activity. We further demonstrated that the MsrPQ repair system ensures Spy’s functionality by reducing these oxidized residues, safeguarding its role in protein homeostasis during oxidative stress. As such, MsrPQ appears to be an essential player in periplasmic proteostasis. Beyond its role in repairing oxidized Met residues, MsrPQ ensures the functionality of critical chaperones like Spy and SurA [12], which collectively prevent protein aggregation and refold damaged proteins [5]. This suggests that MsrPQ likely collaborates closely with chaperones to effectively manage oxidative damage in the periplasm.

Our findings revealed an intriguing paradox: Spy, a chaperone required to mitigate oxidative stress, is itself susceptible to oxidative damage. We demonstrated that mimicking Met oxidation in the disordered N-terminal region has no effect on Spy’s activity. This observation is consistent with the role of the N-terminal region in facilitating the release of client proteins without directly contributing to its chaperone activity [10]. We suspect that the N-terminal Met residues may act as sacrificial “oxidant sponges,” shielding the chaperone’s functional core. This protective role corroborates Stadtman’s theory [27], which states that surface-exposed Met residues shield catalytic sites from oxidative damage. In contrast, the oxidation of Met residues in the protein’s core renders Spy inactive, highlighting their essential role in maintaining its function. The flexibility of Met side chains, due to the presence of sulfur atoms, facilitates protein-protein interactions. These three Met residues likely play a role for structural adaptation during Spy’s interactions with its client proteins. Notably, these Met residues are conserved across all super-variants of Spy obtained from genetic screens aimed at improving chaperone activity against a variety of substrates [25].

Chlorate and chlorite can form naturally in the atmosphere through chlorine photochemistry [28]. However, the primary sources of these compounds are anthropogenic, mainly from the production of bleaching agents and herbicides, leading to environmental contamination. As a result, traces of chlorate and chlorite are found in water and food [29], and could therefore come into contact with the gut microbiota. Our comprehensive proteomic analysis highlights the critical role of Spy and MsrPQ in the bacterial adaptive response to chlorate/chlorite stress. Moreover, it paves the way for future investigations aimed at developing an integrated understanding of this response.

Finally, this work revealed a connection between the oxidative stress response and the envelope stress response in *E. coli*. Indeed, our findings showed that chlorite stress activates not only the HprSR pathway [14], directly linked to oxidative damage, and the BaeSR pathway, traditionally associated with envelope stress [15]. This dual activation suggests an intriguing overlap between these two systems, a link that has received little attention to date. Recent studies in *Salmonella* have similarly observed activation of the Cpx envelope stress pathway by reactive chlorine species such as *N*-chlorotaurine [30], reinforcing the idea of a broader relationship between oxidative stress and envelope integrity.

## MATERIALS AND METHODS

### Construction of strains and plasmids

All strains were derived from *Escherichia coli* MG1655. Lists of strains, plasmids and primers used in this study are shown in Tables 1, 2 and 3 respectively. Gene deletions were performed using the one-step λ Red recombinase chromosomal inactivation system [31]. Deletions were transferred to the WT strain using P1 transduction procedures and verified by polymerase chain reaction (PCR). The antibiotic resistance cassette was removed using Flp recombinases from plasmid pCP20 [31].

**Table 1.**
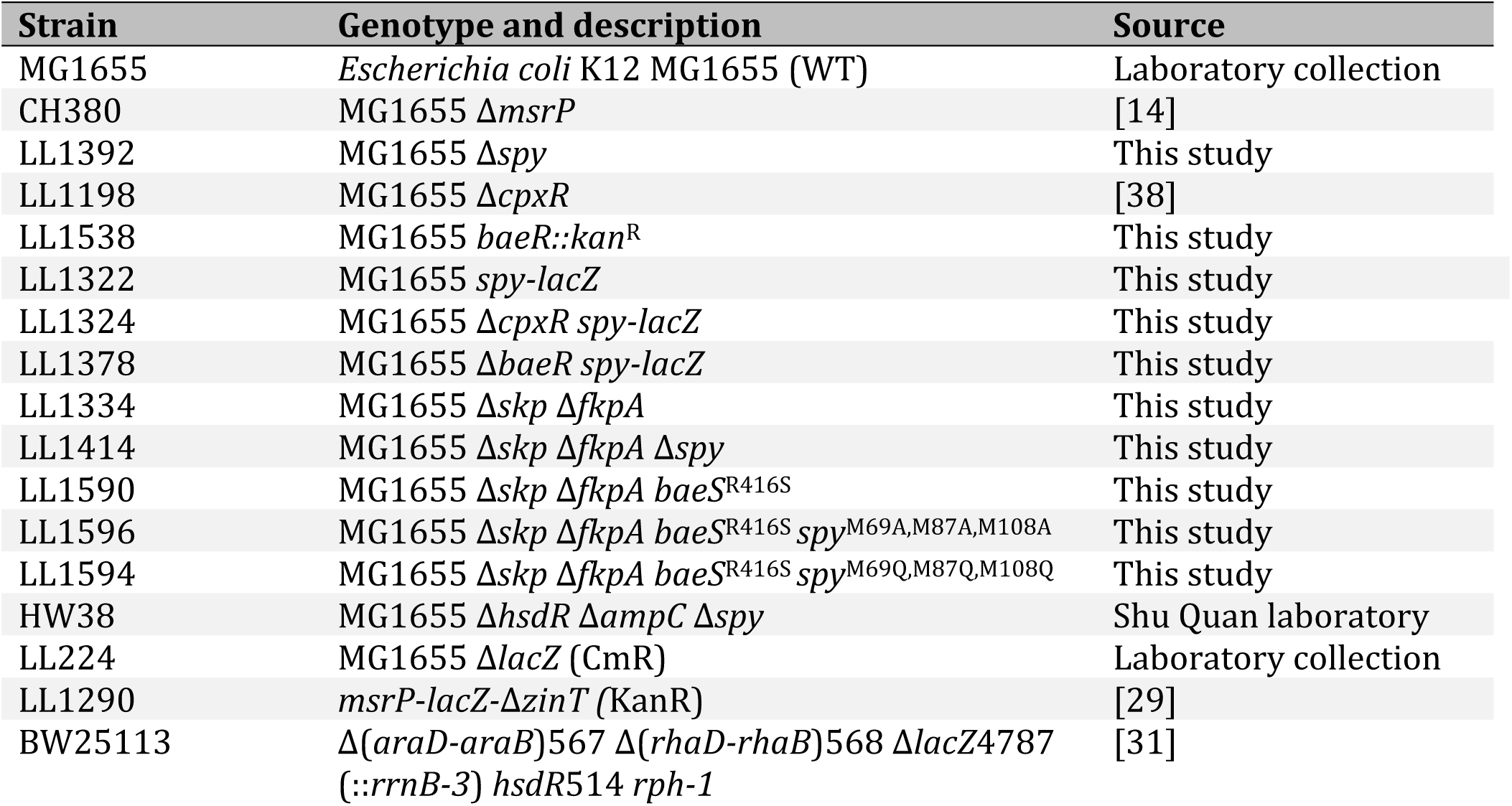
Strains used in this study.

**Table 2.**
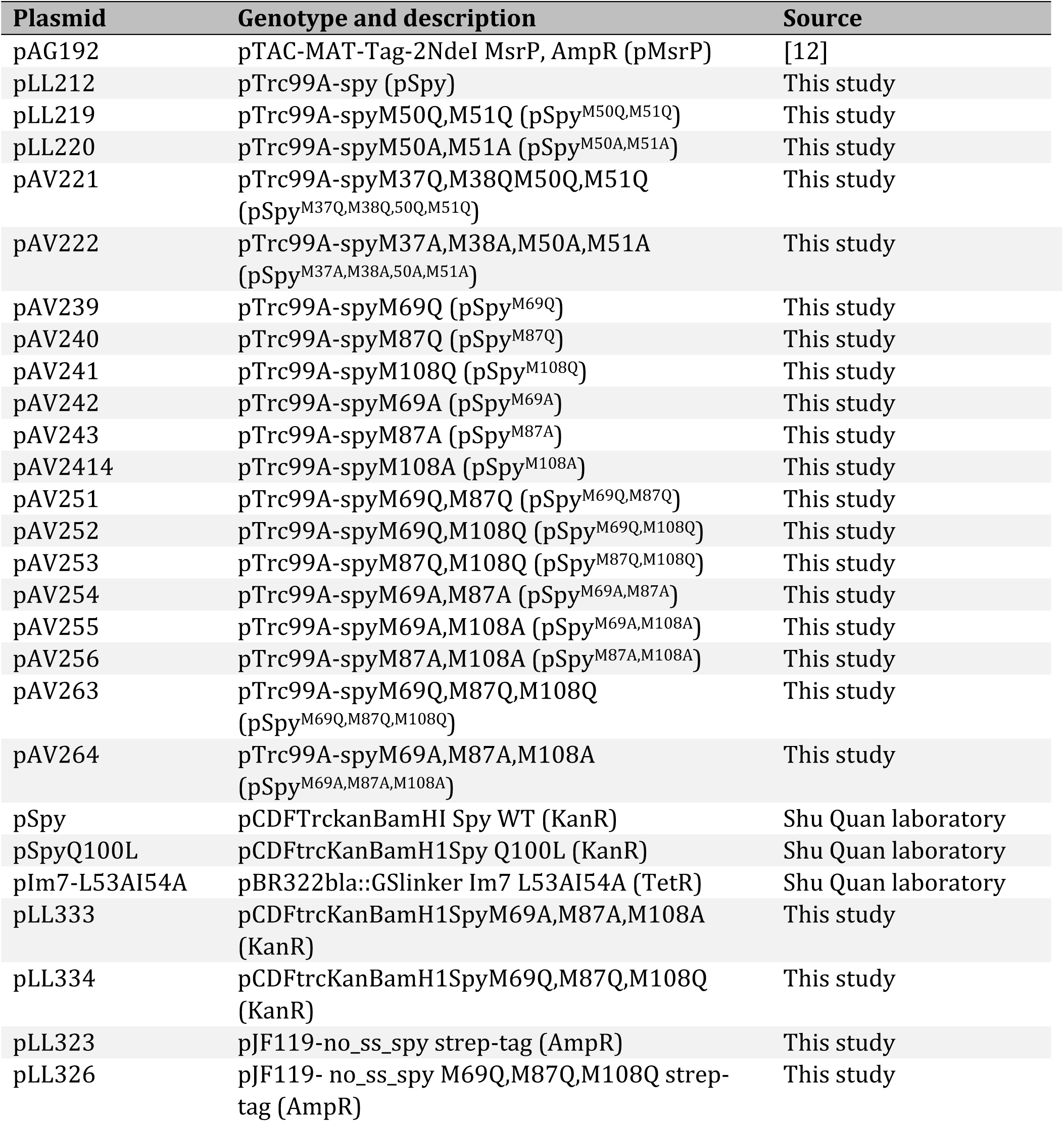
Plasmids used in this study.

| Plasmid | Genotype and description | Source |
| --- | --- | --- |
| pAG192 | pTAC-MAT-Tag-2NdeI MsrP, AmpR (pMsrP) | [12] |
| pLL212 | pTrc99A-spy (pSpy) | This study |
| pLL219 | pTrc99A-spyM50Q,M51Q (pSpy<sup>M50Q,M51Q</sup>) | This study |
| pLL220 | pTrc99A-spyM50A,M51A (pSpy<sup>M50A,M51A</sup>) | This study |
| pAV221 | pTrc99A-spyM37Q,M38Q,M50Q,M51Q<br>(pSpy<sup>M37Q,M38Q,50Q,M51Q</sup>) | This study |
| pAV222 | pTrc99A-spyM37A,M38A,M50A,M51A<br>(pSpy<sup>M37A,M38A,50A,M51A</sup>) | This study |
| pAV239 | pTrc99A-spyM69Q (pSpy<sup>M69Q</sup>) | This study |
| pAV240 | pTrc99A-spyM87Q (pSpy<sup>M87Q</sup>) | This study |
| pAV241 | pTrc99A-spyM108Q (pSpy<sup>M108Q</sup>) | This study |
| pAV242 | pTrc99A-spyM69A (pSpy<sup>M69A</sup>) | This study |
| pAV243 | pTrc99A-spyM87A (pSpy<sup>M87A</sup>) | This study |
| pAV2414 | pTrc99A-spyM108A (pSpy<sup>M108A</sup>) | This study |
| pAV251 | pTrc99A-spyM69Q,M87Q (pSpy<sup>M69Q,M87Q</sup>) | This study |
| pAV252 | pTrc99A-spyM69Q,M108Q (pSpy<sup>M69Q,M108Q</sup>) | This study |
| pAV253 | pTrc99A-spyM87Q,M108Q (pSpy<sup>M87Q,M108Q</sup>) | This study |
| pAV254 | pTrc99A-spyM69A,M87A (pSpy<sup>M69A,M87A</sup>) | This study |
| pAV255 | pTrc99A-spyM69A,M108A (pSpy<sup>M69A,M108A</sup>) | This study |
| pAV256 | pTrc99A-spyM87A,M108A (pSpy<sup>M87A,M108A</sup>) | This study |
| pAV263 | pTrc99A-spyM69Q,M87Q,M108Q<br>(pSpy<sup>M69Q,M87Q,M108Q</sup>) | This study |
| pAV264 | pTrc99A-spyM69A,M87A,M108A<br>(pSpy<sup>M69A,M87A,M108A</sup>) | This study |
| pSpy | pCDFTrckanBamHI Spy WT (KanR) | Shu Quan laboratory |
| pSpyQ100L | pCDFtrcKanBamH1Spy Q100L (KanR) | Shu Quan laboratory |
| pIm7-L53AI54A | pBR322bla::GSlinker Im7 L53AI54A (TetR) | Shu Quan laboratory |
| pLL333 | pCDFtrcKanBamH1SpyM69A,M87A,M108A<br>(KanR) | This study |
| pLL334 | pCDFtrcKanBamH1SpyM69Q,M87Q,M108Q<br>(KanR) | This study |
| pLL323 | pJF119-no_ss_spy strep-tag (AmpR) | This study |
| pLL326 | pJF119- no_ss_spy M69Q,M87Q,M108Q strep-<br>tag (AmpR) | This study |

**Table 3.**
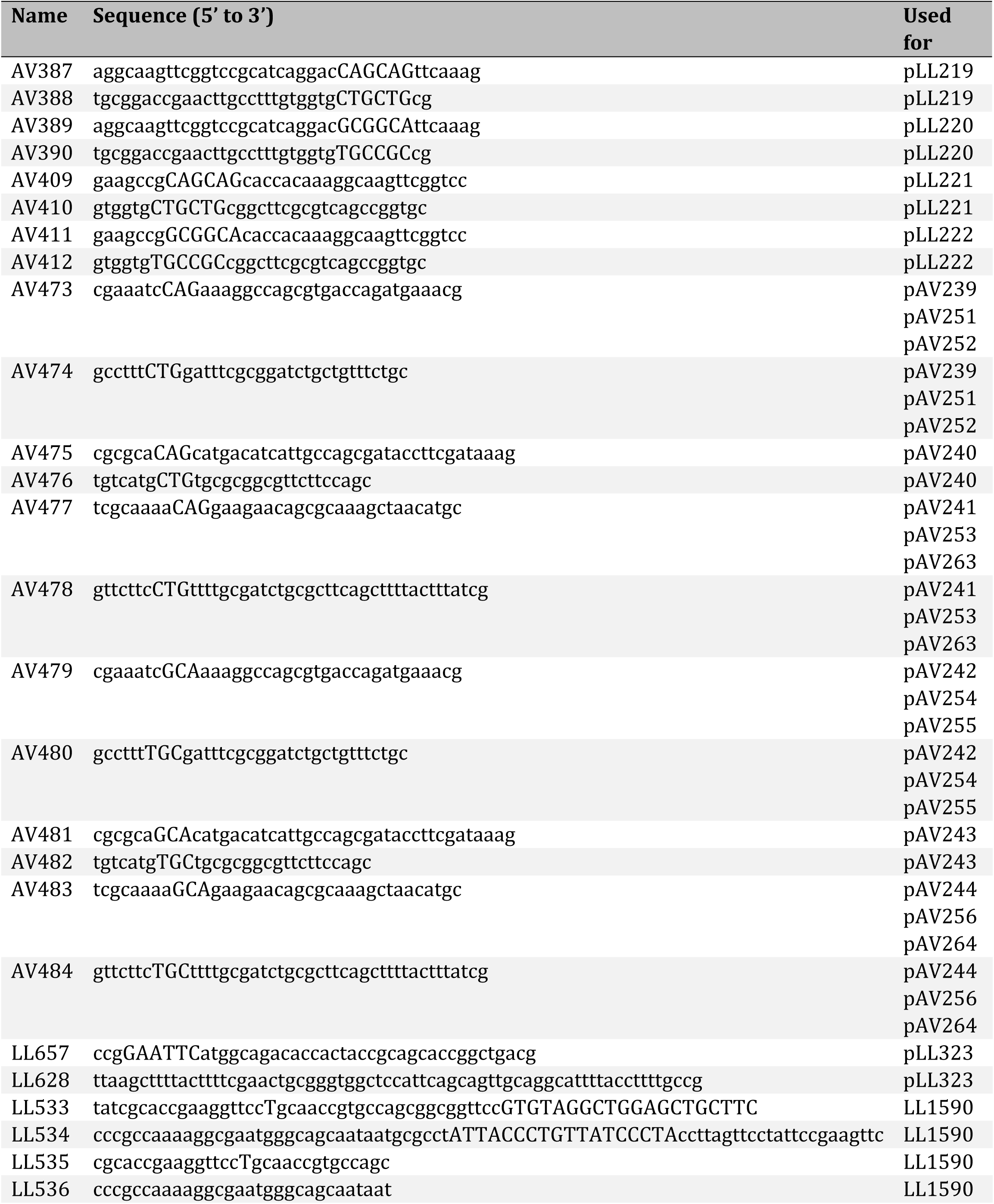
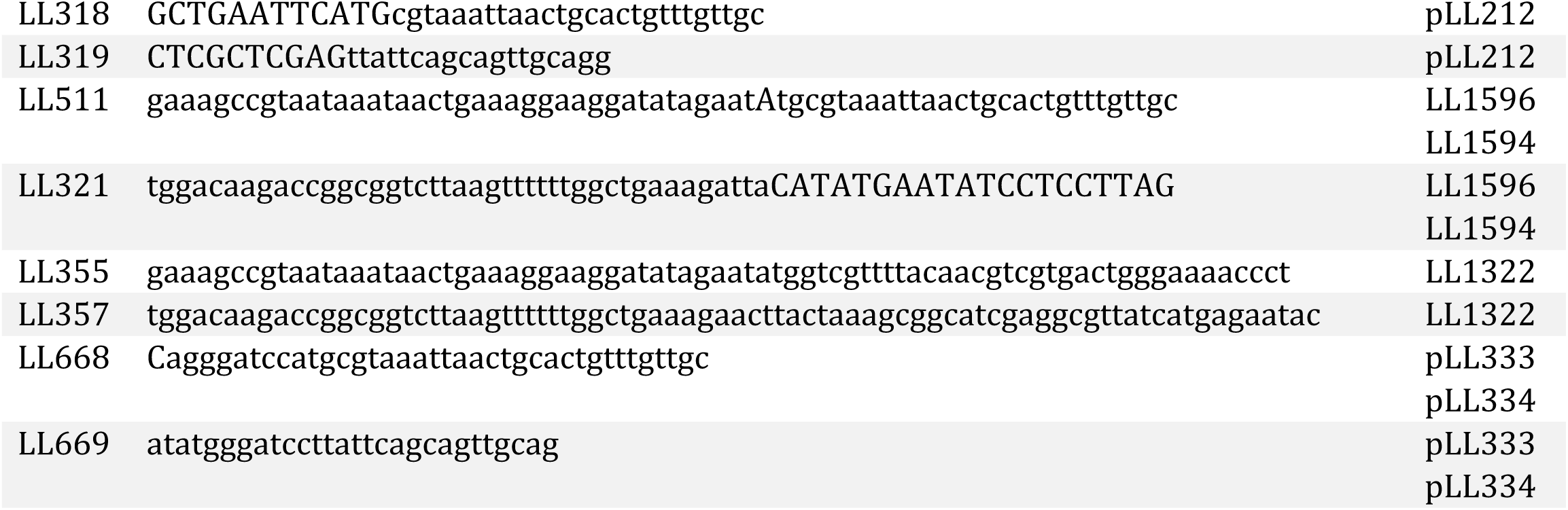
Primers used in this study.

#### Strain engineering

The *spy::lacZ* fusion strain (LL1322) was constructed at the *spy* gene locus. A PCR product containing the *spy* promoter fused to *lacZ* was amplified from strain LL1290 and recombined into BW25113 carrying pKD46. The resulting construct was subsequently transduced into strain LL224 (MG1655 Δ*lacZ*::Cm^R^). The chromosomal *baeS-R416C* strain (LL1590) was generated following procedure described in [32]. The *baeS-R416C sce-I::cat* fragment was amplified by PCR (primers LL533/LL534, pWRG100 template) and electroporated into MG1655 cells expressing λ recombinase from pWRG99. After selection on chloramphenicol (25 µg/mL), the construct was transduced into recipient strains (MG1655, LL1334), and the Cm cassette was removed via λ Red recombination using primers LL535/LL536. The *baeS-R416C* mutation was integrated by electroporation, selected on ampicillin (100 µg/mL) and aTc (1 µg/mL), and confirmed by sequencing. Strains carrying chromosomal *spy* mutant alleles were constructed as follows: plasmids pAV263 and pAV264, containing the mutated versions of *spy*, were opened at the *Hind*III sites. A kanamycin resistance cassette, extracted by *Hind*III digestion from pKD4 vector, was inserted and ligated. These constructs were used as templates for PCR amplification with primers LL511 and LL321 to perform chromosomal insertion by λ Red recombination [36].

#### Plasmid engineering

The IPTG-inducible expression vector for *spy* was constructed by amplifying the *spy* gene from the MG1655 chromosome using primers LL318 and LL319. The PCR product was cloned into pTrc99A via *EcoR*I and *Sal*I restriction sites, generating plasmid pLL212. Plasmids pLL333 and pLL334 (pCDF-Spy^M69A,M87A,M108A^ and pCDF-Spy^M69Q,M87Q,M108Q^, respectively) were constructed by amplifying mutated *spy* alleles from strains LL1596 and LL1594 using primers LL668 and LL669. The PCR products were digested with *BamH*I and cloned into the *BamH*I-linearized pCDFtrcKan vector. For Spy purification, plasmid pLL323 (pJF-no_ss-spy-StrepTag) was constructed by amplifying the *spy* gene (without its signal sequence) from MG1655 using primers LL657 and LL628. This included a Strep-tag II coding sequence at the 3′ end. The PCR product was cloned into pJF119EH using *EcoR*I and *Hind*III restriction sites. The same procedure was used to construct the plasmid pLL326 for expressing Spy^M69Q,M87Q,M108Q^-strepTag, using LL1594 chromosome as the template. All constructs were verified by sequencing.

### Bacterial cultures

*E. coli* strains were streaked onto an LB-agar plate and incubated overnight at 37°C. The following day, a single colony was inoculated into LB and cultured aerobically with shaking at 37°C overnight. The next morning, cultures were diluted to 1:100 prior to performing the experiments described in the following. For immunoblot analyses and β-galactosidase assays, cultures were split into two subcultures and 400 µM ClO_2_⁻ was added to one of the cultures before an additional 4 hours incubation at 37°C with shaking. The following antibiotic concentrations were used for plasmid maintenance: 100 µg.mL^-1^ ampicillin, 25 µg.mL^-1^ chloramphenicol, 30 µg.mL^-1^ kanamycin, 25 µg.mL^-1^ tetracycline.

### Sample Preparation for proteomic analysis

200 µl of the overnight WT culture was diluted into 20 ml of fresh LB medium in a 50 ml Falcon tube. This dilution was split into two 10 ml aliquots in separate 250 ml glass flasks and incubated at 37°C with shaking for 2 hours. A final concentration of 400 µM ClO_2_⁻ was added to one of the flasks. The cultures were incubated at 37°C with shaking for an additional 4 hours. 1 ml of each culture was transferred to pre-weighed 1.5 ml Eppendorf tubes and centrifuged at 8,000 rpm for 2 minutes. The supernatants were removed and the tubes containing the bacterial pellets were weighed again to determine the pellet mass. For every 10 mg of bacterial pellet, 60 µl of Tris-Glycine SDS 2X buffer (Novex) was added. The tubes were heated at 99°C for 5 minutes.

### Shotgun label-free proteomics

Protein extracts (40 µg of total proteins) from three biological replicates under two conditions (cells grown in LB medium, unexposed and exposed to ClO_2_⁻) were subjected to NuPAGE electrophoresis for a short migration of 4 min. The proteomes were treated and subjected to trypsin proteolysis as previously described [33]. The resulting peptides were analyzed on an Orbitrap Exploris 480 (Thermo Scientific) tandem mass spectrometer coupled to a Vanquish Neo UHPLC (Thermo Scientific) and operated as reported [34]. Peptides were desalted on a reverse-phase PepMap 100 C18 μ-precolumn (5 mm, 100 Å, 300 mm i.d. × 5 mm, Thermo Scientific™) and separated on a 50-cm EasySpray column (75 mm, C18 1.9 mm, 100 Å, Thermo Scientific™) at a flow rate of 0.250 μL/min using a 90-min gradient (5–25% B from 0 to 85 min, and 25–40% B from 85 to 90 min) of mobile phase A (0.1% HCOOH/100% H_2_O) and phase B (0.1% HCOOH/100% CH_3_CN). Tandem mass spectrometry results were interpreted as recommended [35].

### Immunoblot analysis

Similar cell densities (10 units/ml; normalized by *A*_600_) were pelleted by centrifugation (3000 g, 5 min), resuspended in Laemmli SDS buffer, heated for 10 min at 95°C, and separated by gel electrophoresis (Novex 4-20% Tris-glycine Gel (ThermoFisher Scientific)). Samples were transferred to 0.2 µm nitrocellulose membranes and probed with anti-Spy (Clinisciences) and anti-MsrP antibody (provided by the Jean-François Collet Lab, de Duve Institute, UCLouvain), followed by an HRP-conjugated anti-Rabbit (Promega W4011) for Spy and anti-guinea pig IgG secondary antibody (Sigma-Aldrich) for MsrP. Chemiluminescence signal was collected using an ImageQuant Las4000 camera (GE Healthcare). The presented results are representative of at least three independent experiments.

### ß-galactosidase assays

100 µl of culture treated or untreated with 400 µM ClO_2_⁻ were added to 900 µl of β-galactosidase buffer. Levels of ß-galactosidase were measured as previously described [36]. The same protocol was employed to assess the effect of other oxidants on *spy* expression. The oxidants were added to LB medium to achieve final concentrations of 5 mM H_2_O_2_ (Honeywell), 0.3 mM paraquat (Sigma-Aldrich), 0.3 mM diamide (Sigma-Aldrich), 4 mM HOCl (Honeywell), and 1 mM *N*-chlorotaurine. 0.25 mM tannic acid (Sigma-Aldrich) was used as a control.

### Sphaeroplasts preparation

A 2 ml sample was centrifuge, washed and resuspended in 250 µl of Tris-HCl 0.2 M, pH 8 and mixed with 250 µl of Tris-HCl 0.2 M, pH 8, sucrose 1 M, EDTA 1 mM. 2 µl of lysozyme (15 mg/ml) was added and the suspension was incubated at room temperature for 30 minutes. The suspension was centrifuged, and the supernatant was harvested to collect the periplasmic fraction.

### *N*-ChT synthesis

*N*-ChT (Cl-HN-CH_2_-CH_2_-SO_3_^-^) was produced by mixing 100 mM HOCl (Honeywell) and 100 mM taurine (Sigma-Aldrich) in 0.1 M phosphate potassium buffer (pH 7.4). After 10 min at room temperature, the *N*-ChT concentration was determined by measuring the absorbance at 252 nm (ɛ = 429 M^-1^.cm^-1^) and stored at 4°C.

### Site-directed mutagenesis

50 µl PCR reactions were performed using Q5 Hot start High-Fidelity DNA polymerase (New England Biolabs). The resulting PCR products were digested with *Dpn*I, purified using GeneJET PCR purification kit (Thermo Fisher) and transformed into DH5α. Three colonies were randomly selected from each transformation, and the plasmids were purified using GeneJET Plasmid Miniprep kit (Thermo Fisher) and verified by sequencing. Templates and primers used for generating Spy variants are shown in Table. 4.

**Table 4.** Templates et primers used to construct *spy* variants.

| Templates | Primers | Products |
| --- | --- | --- |
| pLL212 | AV387/AV388 | pLL219 |
| pLL212 | AV389/AV390 | pLL220 |
| pLL219 | AV409/AV410 | pLL221 |
| pLL220 | AV411/AV412 | pLL222 |
| pLL212 | AV473/AV474 | pAV239 |
| pLL212 | AV475/AV476 | pAV240 |
| pLL212 | AV477/AV478 | pAV241 |
| pLL212 | AV479/AV480 | pAV242 |
| pLL212 | AV481/AV482 | pAV243 |
| pLL212 | AV483/AV484 | pAV244 |
| pAV240 | AV473/AV474 | pAV251 |
| pAV241 | AV473/AV474 | pAV252 |
| pAV240 | AV477/AV478 | pAV253 |
| pAV243 | AV479/AV480 | pAV254 |
| pAV244 | AV479/AV480 | pAV255 |
| pAV243 | AV483/AV484 | pAV256 |
| pAV251 | AV477/AV478 | pAV263 |
| pAV254 | AV483/AV484 | pAV266 |

### Novobiocin and clindamycin survival assays

The Δ*skp* Δ*fkpA* Δ*spy* strain (LL1414) harboring an empty vector, pSpy wild type or mutated variants grew aerobically in LB supplemented with ampicillin and IPTG (0.1 mM) at 37°C. At OD_600_ = 1, cultures were serially diluted in phosphate buffered saline (PBS). 3.5 µl of 10-time serial dilutions were spotted onto LB-agar plates containing ampicillin and IPTG (0.1 mM) supplemented or not with 50 µg/ml novobiocin or 75 µg/ml clindamycin. Plates were incubated at 37°C overnight. For the clindamycin sensitivity experiments without plasmid addition, the protocol was identical, except ampicillin and IPTG were omitted. LB plates supplemented with 50 µg/mL clindamycin were used.

### Chaperone activity assay *in vivo*

Spot titer experiments were performed to quantify the relative *in vivo* chaperone activity of Spy variants, as previously established and described [37]. Briefly, Δ*hsdR* Δ*ampC* Δ*spy* cells (HW38) expressing Spy variants from a pCDFTrckanBamHI and ßla-Im7 L53A I54A sensor from pBR322 plasmids (provided by the Shu Quan Lab, Shanghai, Jiao Tong University) were grown in LB supplemented with kanamycin, tetracycline and IPTG (0.1 mM) to OD_600_ = 0.6, serially diluted, and plated onto LB-agar plates supplemented or not with penicillin V (1 and 1.5 mg/ml) and IPTG (0.1 mM). Plates were incubated at 37°C overnight.

### Purification of Spy

Strep-tagged versions of wild-type Spy and its variants were expressed and purified from *Δspy* (LL1392) cells harboring plasmids pLL323 or pLL326, which overexpressed Spy^WT^ or Spy^M69Q,M87Q,M108Q^ proteins, respectively. Cells were grown aerobically at 37°C in LB medium supplemented with ampicillin. When cultures reached OD_600_ = 0.6, protein expression was induced with IPTG (0.1 mM final concentration) for 4 hours at 37°C. The pellet from a 400 ml culture was resuspended in buffer A (100 mM Tris-HCl, 50 mM NaCl, pH 8). Cells were disrupted by two passes through a French press. After centrifugation at 12,000 rpm for 30 minutes at 4°C to remove debris, the supernatant was loaded onto a 5- ml Strep-Trap HP column (GE Healthcare) equilibrated with buffer A. The column was washed with buffer A, and Spy was eluted using buffer A supplemented with 2.5 mM desthiobiotin. Pure fractions were pooled, and desthiobiotin was removed using an Amicon Ultra-10K filter (Millipore).

### α-lactalbumin aggregation assay

Bovine α-lactalbumin aggregation assay was performed as previously described [10,25,26]. Briefly, aggregation of α-LA (type III,Sigma Aldrich) at 100 µM was initiated by adding 20 mM DTT to a buffer containing 50 mM phosphate buffer, 100 mM sodium chloride, and 5 mM EDTA, pH 7.0. The light scattering of α-LA in the absence or presence of Spy (30 µM) was monitored at 360 nm using a Spark microplate reader (TECAN) with a 5-min detection period for 600 min at 25 °C.

## ACKNOWLEDGEMENTS

We thank members of the Ezraty group for their discussions. We are grateful to James Bardwell (University of Michigan), Shu Quan (Shanghai Jiao Tong University) and Jean-François Collet (de Duve Institute, UCLouvain) for their advice, discussions, and for providing strains and plasmids. We also thank Marianne Ilbert and Olivier Genest (BIP-AMU/CNRS Marseille) for their valuable advice, as well as Mélodie Kielbasa (CEA-Li2D) for her technical help with mass spectrometry. This work was supported by grants from the Agence Nationale Recherche (ANR) (#ANR-16-CE11-0012-02 METOXIC). J.A. acknowledges the France2030 program – INBS ProFI (grant ANR-24-INBS-0015). M.S.V was funded by the European Union’s Horizon 2022 programme under a Marie Skłodowska-Curie MSCA postdoctoral fellowship No 101106503 (MOR-AGE). This work received support from the French government under the France 2030 investment plan, as part of the Initiative d’Excellence d’Aix-Marseille Université - A*MIDEX and is part of the Institute of Microbiology, Bioenergies and Biotechnology - IM2B (AO-IM2B-NE-2024-02-VINCENT).

The authors declare no conflict of interest.

